# Quantile-specific confounding: correction for subtle population stratification via quantile regression

**DOI:** 10.1101/2025.03.18.638253

**Authors:** Chen Wang, Marco Masala, Edoardo Fiorillo, Marcella Devoto, Francesco Cucca, Iuliana Ionita-Laza

## Abstract

Subtle population structure remains a significant concern in genome-wide association studies. Using human height as an example, we show how quantile regression, a natural extension of linear regression, can better correct for subtle population structure due to its inherent ability to adjust for quantile-specific effects of covariates such as principal components. We utilize data from the UK biobank and the SardiNIA/ProgeNIA project for demonstration.

---

Many complex traits are highly polygenic, and genome-wide association studies (GWAS) allow for the discovery of genetic variation associated with such traits. A well-known confounder in GWAS is population structure; left unaccounted, it leads to false positive associations. Adjusting for this confounding effect using principal component analyses (PCA) or mixed effect modeling has become standard practice in GWAS [1]; yet there is always the risk of residual confounding especially in large and heterogeneous GWAS analyses. Indeed, recent studies have shown that population stratification still accounts for a substantial fraction of GWAS associations for many traits [2], including height. The largest GWAS for height to date with approximately 5.4 million individuals has been conducted by the GIANT (Genetic Investigation of Anthropometric traits) consortium [3]. GIANT is by nature a large and heterogeneous dataset that includes a vast number of cohorts of mostly European-descent individuals from Europe and North America, and more than one million participants of East Asian, Hispanic, African, or South Asian ancestry, and therefore concerns about incomplete control of population structure have been raised in the literature [4–5]. Even in the UK Biobank (UKBB), a much more homogeneous cohort, there is evidence of residual population stratification [6].

PCA performs well when the structure has a smooth and wide distribution. However, for more sharp and localized distributions, PCA may not be enough [7]. Although alternative methods for detecting finer scale population structure have been discussed in the literature [8,9], there are several drawbacks and practical challenges for implementing them in practice.

We illustrate here an application of quantile regression (QR) [10], a classical statistical technique that can better adjust for sharp and localized population stratification. In such cases, the confounding effect may vary across parts of the phenotype distribution and may even be concentrated on specific quantiles. Linear regression (LR) by design assumes that covariate effects are homogeneous across quantiles and therefore can under- or over-adjust when effects are heterogeneous. In contrast, QR allows for covariate adjustment at specific quantiles, and therefore can be more appropriate than LR when the covariate effects vary across quantiles. For example, physical activity can have stronger effects on upper BMI levels, but lower effects on upper height quantiles ([11], **Fig. S1**) and QR naturally adjusts for these heterogeneous effects.

QR is a natural extension of LR and consists of fitting a series of linear regression models at different quantile levels *τ* ∈ (0,1). To test for association between the *j*th genetic variant and phenotype *Y* at quantile level *τ* we fit the following conditional quantile regression model:

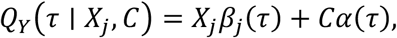

where *Q*_*Y*_(*τ*|*X, C*) is the *τ*th conditional quantile of phenotype *Y* given genotypes *X* and covariates *C*; *β*_*j*_(*τ*) and *α*(*τ*) are quantile-specific coefficients and can be estimated by minimizing the pinball loss function [10]. Since both the genotype effects *β*_*j*_(*τ*) and the covariate effects *α*(*τ*) are allowed to vary by quantile, QR can (*i*) detect heterogeneity in effects across quantiles, and (*ii*) adjust for covariates with varying effects across quantiles. For testing the null hypothesis *H*_0_: *β*_*j*_(*τ*) = 0, we employ the rank score test to test and obtain a p-value for each SNP at quantile level *τ* [12–13]. In practice, in addition to quantile specific p-values we also compute a composite p-value by combining p-values at multiple quantile levels using Cauchy’s combination method [14].

To illustrate the scenario of local and sharp population stratification, we utilized data from 325,667 white British unrelated individuals in UKBB, and 1,946 unrelated individuals from the ProgeNIA/SardiNIA project, a longitudinal study of a cohort of Sardinian subjects [15–16]. Following standard QC procedures, we analyzed 7,844,680 SNPs in both datasets. We used standing height measurements (at baseline) as the phenotype.

We performed LR as well as QR at several quantile levels, including *τ* = 0.1, 0.3, 0.5, 0.7, 0.9. For both LR and QR, we adjusted for sex and 10 global PCs (**Methods**). We did not adjust for age, as age can act as a collider in the combined Sardinia and UKBB data leading to high inflation in false positives.

British individuals are on average taller than individuals from Sardinia by 8 cm (**Fig. 1(a)**), leading to more pronounced stratification at lower quantile levels (e.g. *τ* = 0.1 − 0.3). PC1 and PC2 are describing genetic variation within UK, while PC3 and PC5 can separate the two subpopulations (**Fig. 1(b)**). The effects of PC5 on height vary as a function of quantile levels and are stronger at lower quantiles (**Fig. 1(c)**).

**Figure 1:**
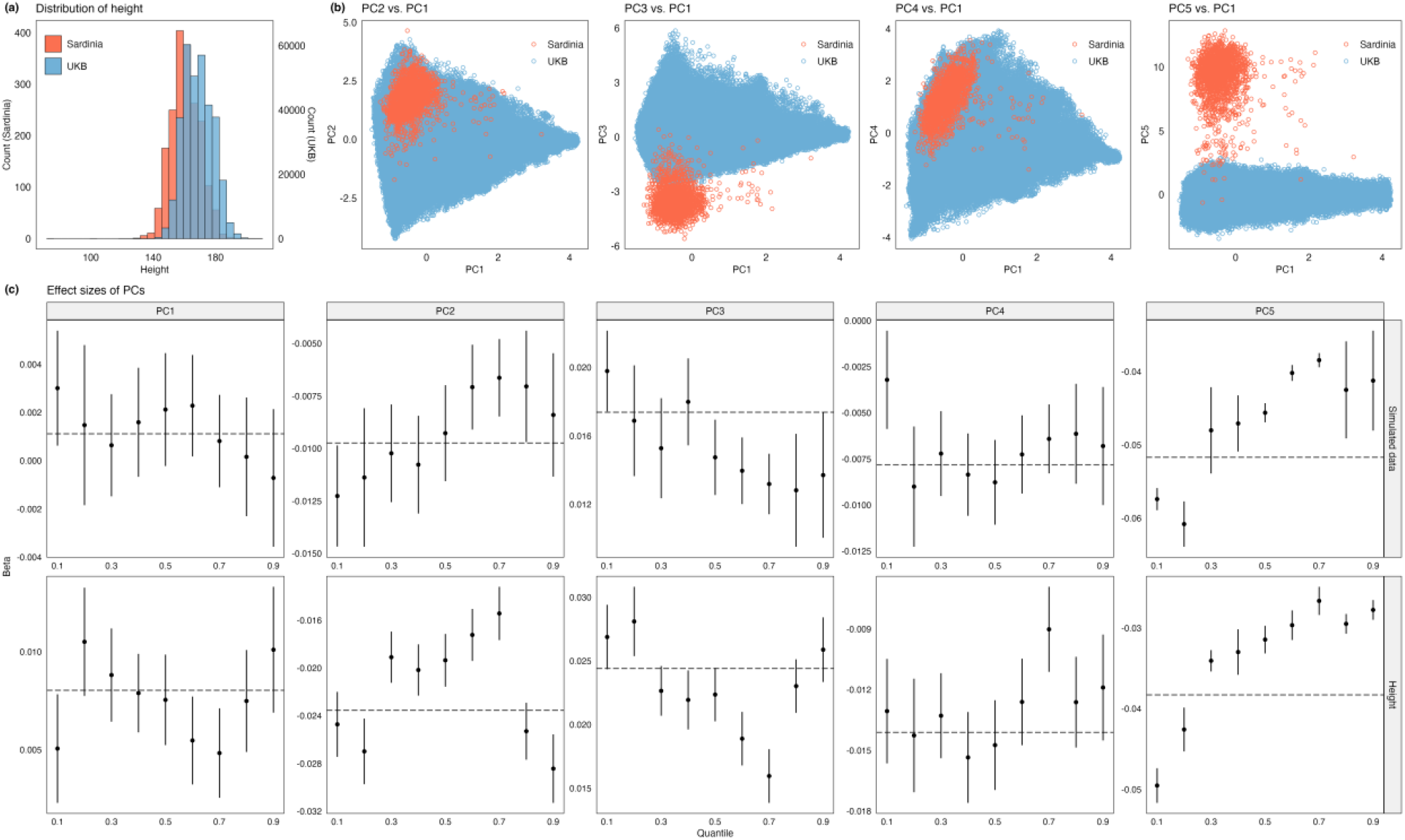
Height distribution and PCs of genetic variation in two cohorts, Sardinia and UKBB. **(a)** Height distributions; **(b)** PC plots for top PCs; **(c)** PCs’ estimated quantile-specific effects (and standard errors) on height at quantile levels *τ* = 0.1 − 0.9. Upper panel: simulated data; lower panel: height. The horizontal line corresponds to the mean-based effect in LR.

We first performed simulations by randomly permuting the height phenotype separately within the Sardinia and UKBB data, and report detailed results for one replicate and summary results across 20 independent replicates. We noticed a decrease in false positives induced by population stratification for QR relative to LR (**Fig. 2(a)**). Specifically, LR tends to identify more significant associations than QR across different replicates.

**Figure 2:**
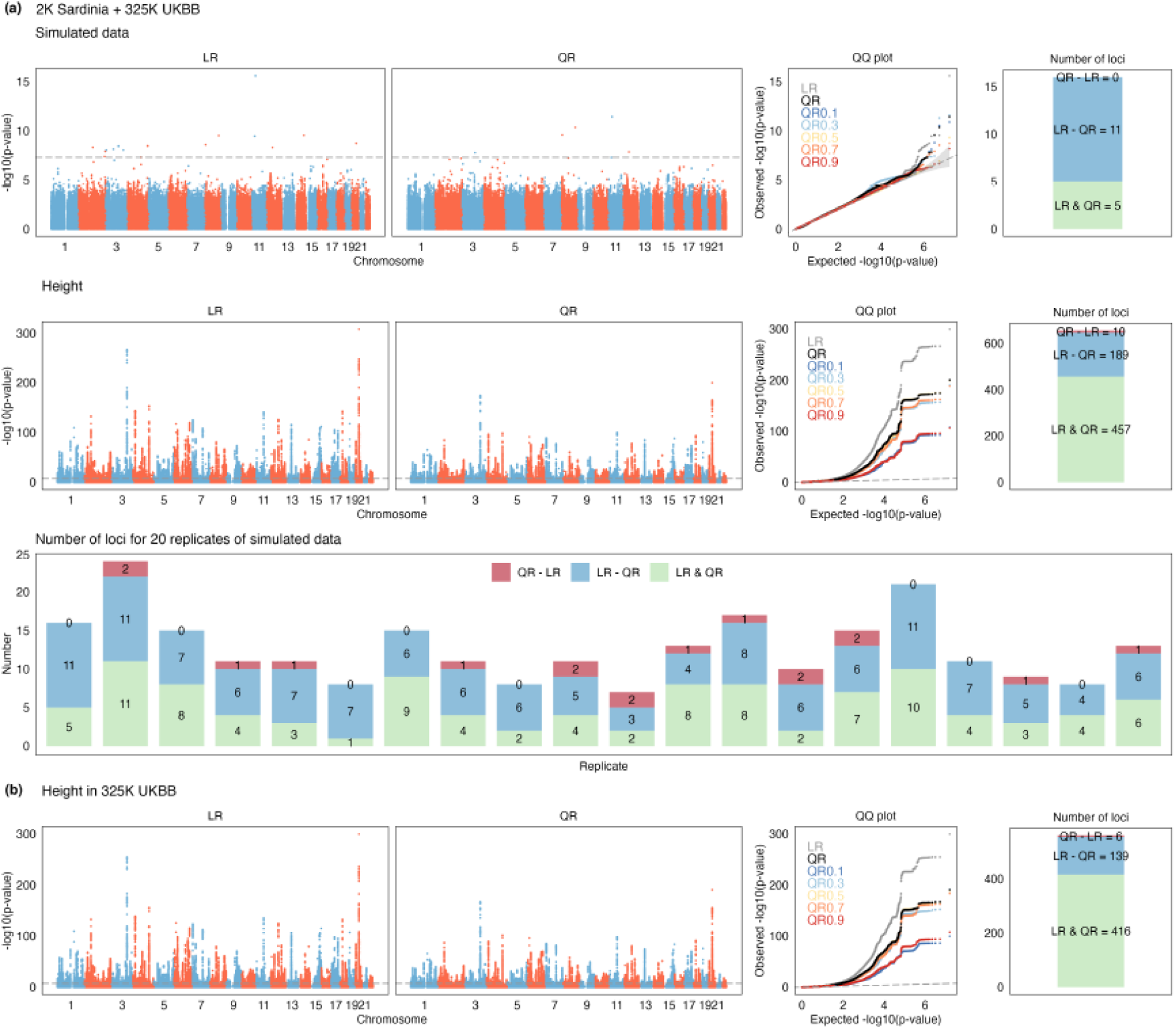
LR and QR GWAS on simulated and height data with stratification. Manhattan and QQ plots are shown for **(a)** simulated (no real signal) and height data when combining UKBB and Sardinia; **(b)** only UKBB. We report the number of independent loci (unique and shared) identified by LR and QR.

Similarly, for real height, we see a reduction in the number of significant associations for QR vs. LR (**Fig. 2(a)**). Although most associations are shared between LR and QR (457 loci), LR results in a much larger number of new discoveries relative to QR (189 vs. 10 loci). These new associations likely reflect both increased power for LR over QR but also increased number of false positives since by combining data from UKBB and Sardinia a certain number of false positives are expected due to effects of population stratification and assortative mating. When restricting analyses to 325,667 white British unrelated individuals, the number of discoveries decreases substantially, especially for loci identified by LR but missed by QR (139 vs. 189 above) likely reflecting an overall reduction in false positives (**Fig. 2(b)**).

By comparison, in the case of BMI the effect of population stratification appears less pronounced (**Fig. S2** and **S3**). Relative to height, BMI shows more heterogeneity in genetic effects genome-wide, with power at the upper quantiles higher than at lower quantiles, suggestive of more pronounced gene-by-environment effects at upper quantiles of BMI, as expected.

In summary, QR is a robust technique for the analysis of continuous phenotypes in large heterogeneous genomic data that provides a good balance between discovery power and control of false positives. Beyond its robustness, QR allows for a more nuanced understanding of genotype-phenotype correlations by revealing patterns of heterogeneity that is not possible with the GWAS approach.

## Acknowledgements

C.W. and I.I.L. are supported by NIH grants MH095797 and AG072272. This research has been conducted using the UK Biobank Resource under Application Number 41849.

## Code Availability

Scripts for QR GWAS analysis are available at https://github.com/Iuliana-Ionita-Laza/QRGWAS. The quantile regression was performed using the R package quantreg (https://cran.r-project.org/package=quantreg) version 5.94. The rank score test was performed using the R package QRank (https://cran.r-project.org/package=QRank) version 1.0.

## Methods

### Overview of quantile regression

We assume that we have n independent samples from a population. We denote by Y = (Y_1_, ⋯, Y_n_)^′^ the n × 1 sample phenotype vector, by X the n × p genotype matrix, and by C the n × q matrix for covariates including age, gender and principal components of genetic variation. In GWAS, the typical approach is to perform marginal (unconditional) testing for one variant at a time, assuming the following model:

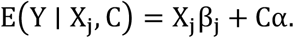

In QR, we denote the *τ*th conditional quantile function of Y as Q_Y_(τ ∣ X, C). For linear QR, we can write the conditional quantile regression model for the jth variant and specific quantile level τ ∈ (0,1) as:

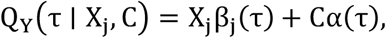

where β_j_(τ) and α(τ) are quantile-specific coefficients that can be estimated by minimizing the pinball loss function. This optimization problem can be formulated equivalently as a linear programming problem that can be solved efficiently using the simplex algorithm or interior point methods. For QR, a commonly used hypothesis testing tool is the rank score test that allows us to obtain well-calibrated p-values at individual quantile levels. The rank score test statistic for a fixed quantile level *τ* is defined as:

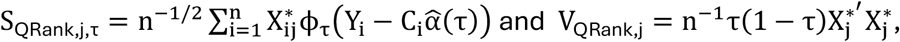

where X^*^ = P_C_X, P_C_ = I − C(C^′^C)^−1^C′ and Φ_*τ*_(u) = τ − I(u < 0) is the τ-pinball loss. Under the null hypothesis 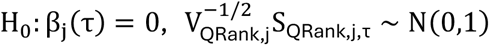. Note that the asymptotic distribution of the test statistic is independent of the distribution of the phenotype. Hence it can be applied to any phenotype without requiring a pre-transformation to achieve normality, which is another important advantage of QR over LR. Cauchy combination can be used to integrate results over different quantile levels.

### PCA

We perform PCA analysis as follows. We combined genotype data of unrelated samples from the UKBB White British population (n=325,667) and the ProgeNIA/SardiNIA cohort (n=1,946). To identify unrelated individuals, we used KING [1] to estimate the genetic relatedness between samples and only retained the unrelated samples that were more distant than 2nd-degree relatives (KING kinship coefficient <= 0.0884). We selected SNPs with a minor allele frequency (MAF) > 0.01 and genotype missingness < 5%. We performed linkage disequilibrium (LD) pruning using PLINK2 [2] with a sliding window of 1000 SNPs, a step size of 100 SNPs, and an LD threshold r2 < 0.1 to retain independent variants. We also excluded three long-range LD regions (chr6: 25.5M-33.5M, chr8: 8M-12M, and chr11: 46M-57M) [3] from the LD pruning. We computed the first 10 PCs on the LD-pruned variants using randomized partial PCA implemented in FlashPCA [4].

### Locus identification

We performed LD clumping using PLINK2 [2] with a 1 Mb distance threshold and an LD threshold of r2 > 0.1 to identify independent loci. Adjacent loci within 1 Mb of each other were subsequently merged into a single locus.

---

**Figure S1:**
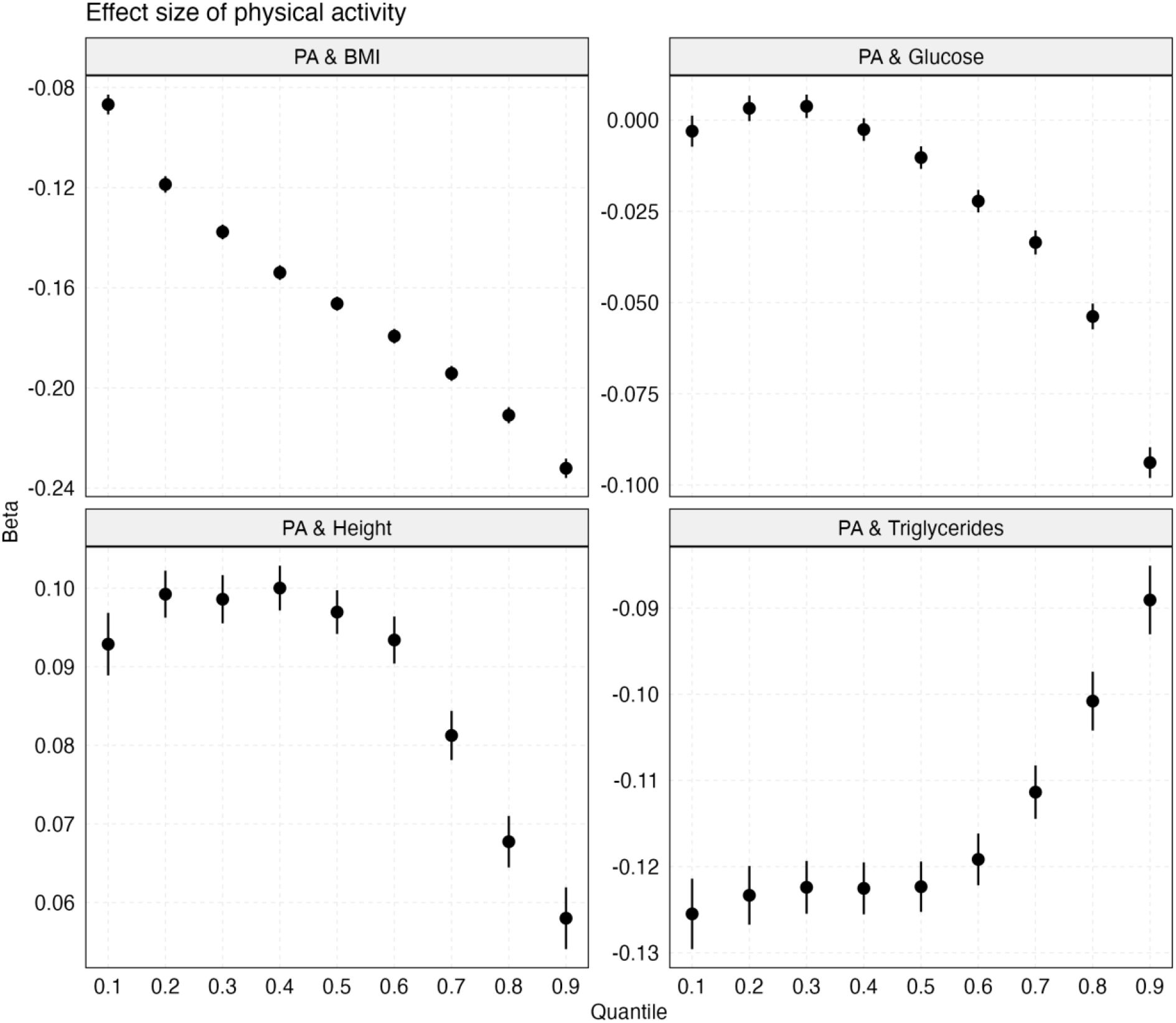
Heterogeneous effects of physical activity (PA) on BMI, height, glucose and triglycerides in UKBB. Estimated effects and standard errors across different quantile levels are shown. These analyses are based on unrelated white British individuals (between 264,854 and 325,825 depending on the trait). PA is a categorical variable (high, moderate, low) defined based on questionnaire data.

**Figure S2:**
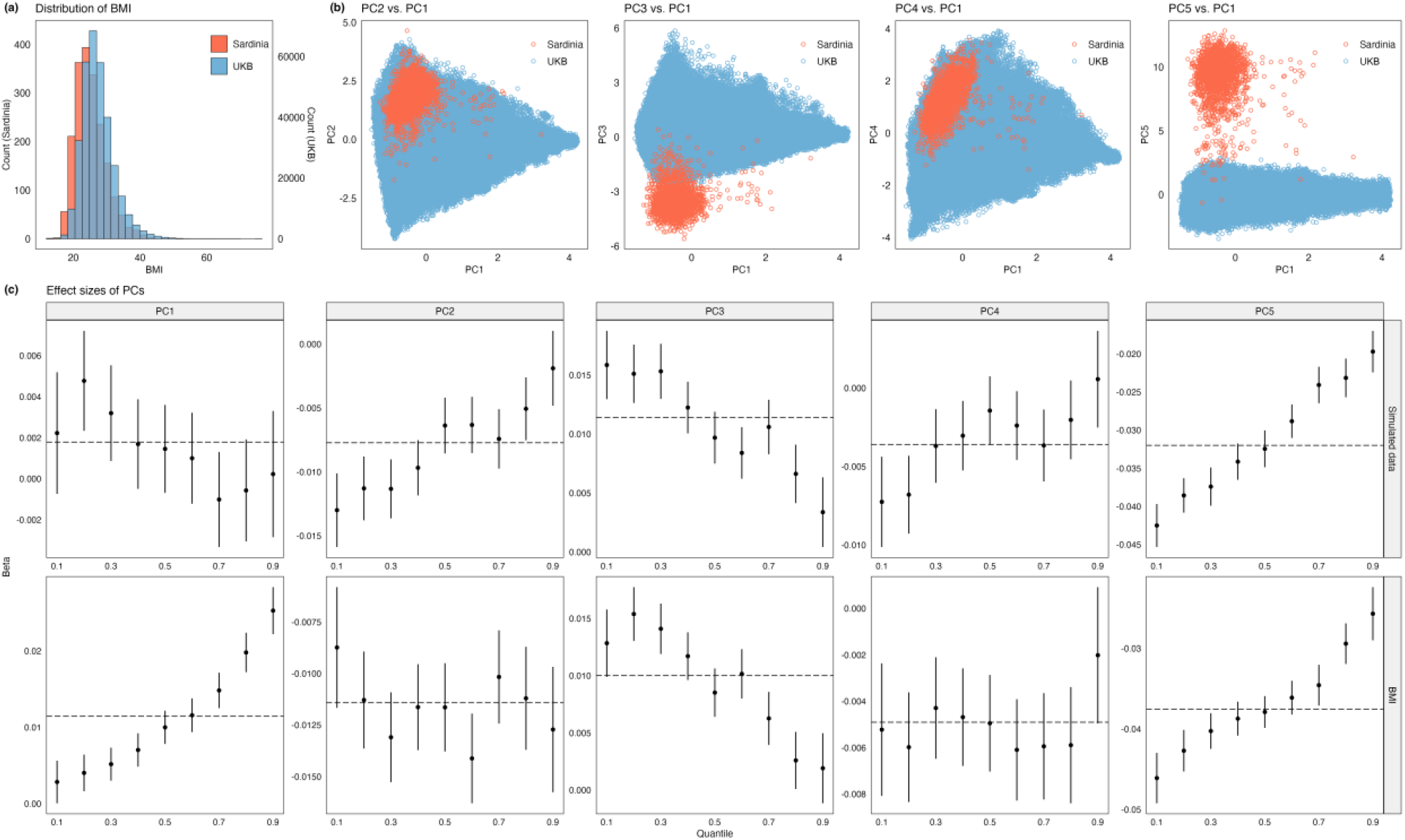
BMI distribution and PCs of genetic variation in two cohorts, Sardinia and UKBB. **(a)** BMI distributions; **(b)** PC plots for top PCs; **(c)** PCs’ estimated quantile-specific effects (and standard errors) on BMI at quantile levels *τ* = 0.1 − 0.9. Upper panel: simulated data; lower panel: BMI. The horizontal line corresponds to the mean-based effect in LR.

**Figure S3:**
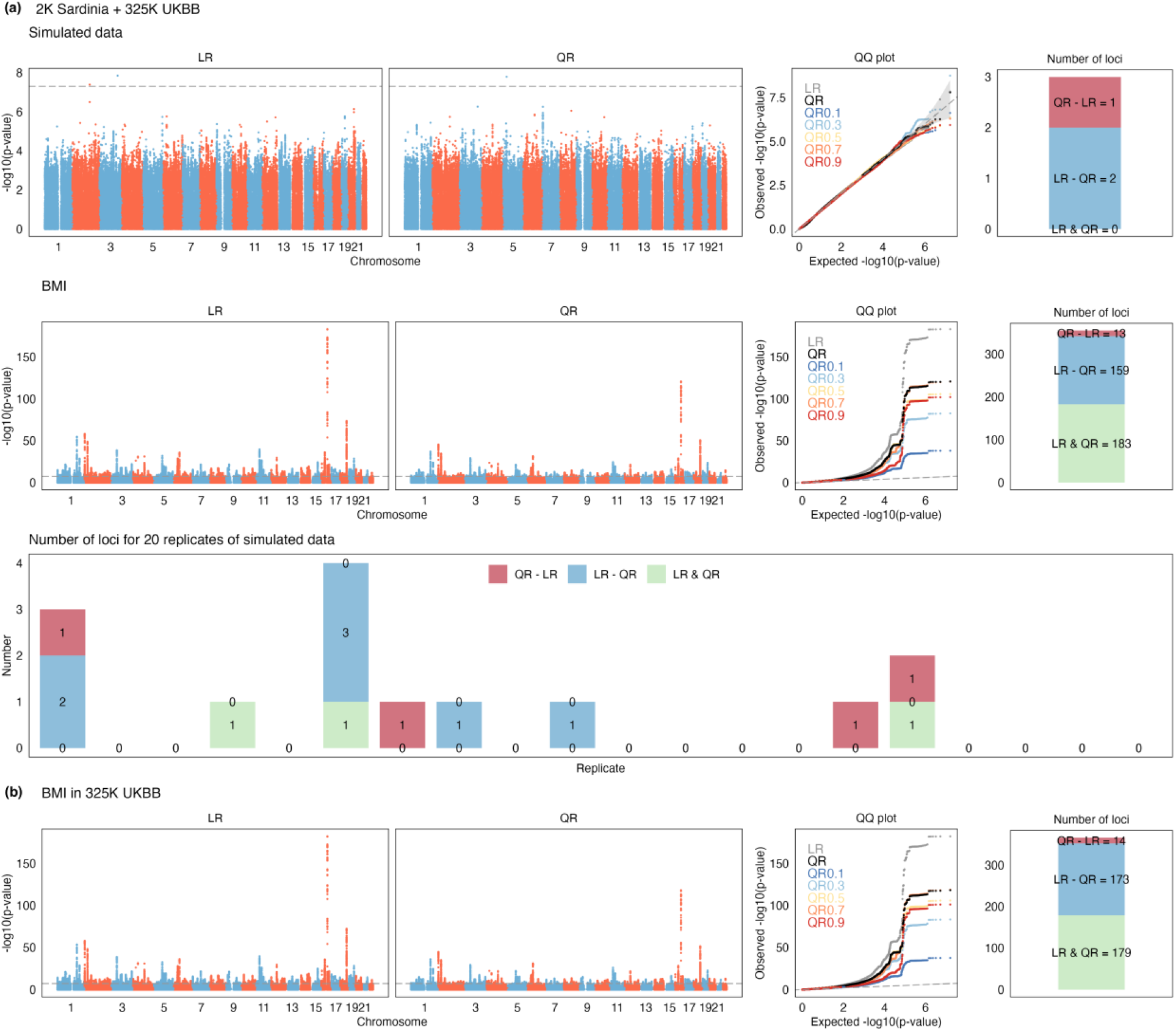
LR and QR GWAS on simulated and BMI data with stratification. Manhattan and QQ plots are shown for **(a)** simulated (no real signal) and BMI data when combining UKBB and Sardinia; **(b)** only UKBB. We report the number of independent loci (unique and shared) identified by LR and QR.

